# Flight heights of Scopoli’s shearwaters *Calonectris diomedea* in the context of offshore wind farm developments

**DOI:** 10.1101/2023.05.14.540698

**Authors:** Nicolas Courbin, Aurélien Besnard, Etienne Boncourt, David Grémillet

## Abstract

Seabird face new risks through collisions with offshore wind turbines. Wind energy projects are emerging in the Mediterranean Sea, and unbiased knowledge on seabird flight altitudes are scarce in this area. Indeed, previous flight height observations were carried out from boats during daytime, and only in good weather conditions. We measured flight heights of 13 Scopoli’s shearwaters, *Calonectris diomedea*, one of the three endemic shearwater species in the Mediterranean, using highly precise barometric data loggers. The birds were equipped at the largest French colony of Scopoli’s shearwater in the Marseille archipelago. Birds from this location routinely visit forthcoming wind farm areas, within 50 km of their breeding site. We found that Scopoli’s shearwaters flew at very low altitudes with a mean flight height of 1.8 ± 2.7 m (± SD) above sea level. Birds therefore rarely (<0.02%) flew within the vertical envelop of collision risk with wind turbines. Ours results are coherent with field observations and the dynamic soaring flight technique used by shearwaters.

Nevertheless, collision risk may increase following behavioural modifications close to wind turbines.

## INTRODUCTION

Worldwide seabird populations suffered a 50% decline between 1970 and 2010 (Grémillet et al., 2018) and seabirds are, together with parrots, the most vulnerable bird group (Dias et al., 2019, Rodríguez et al. 2019). The accumulation of anthropogenic stressors on adult survival is of primary concern considering the key role of adult survival for the population dynamic of these long-lived species, characterized by a slow pace of life (Saether & Bakke, 2000; Caswell, 2001). Seabird face to a new risk through collisions with offshore wind turbines.

Indeed, the current race towards decarbonised energy leads a huge development of wind energy, with a considerable increase in the potential spatial extent for offshore wind farms worldwide (International Energy Agency, 2019). The total installed offshore wind power capacity in the European Union (currently ca. 200 GW) might be multiplied by seven until 2040 (International Energy Agency, 2019). Offshore wind farms have substantial ecological impacts on seabirds (Furness et al. 2013; Green et al., 2016; Galparsoro et al., 2022), and the nature and magnitude of their effects vary considerably across seabird species (Bradbury et al., 2014; Thaxter et al., 2017). Considering the precarious situation of most seabird populations, even modest additional individual mortalities from wind turbines can have strong detrimental effects on populations. Quantifying the potential effect of collisions with turbines on seabird population dynamics is nonetheless challenging, since it requires the combined analysis of precise data on flight altitude and space use of birds, population sizes and demographic parameters (Cleasby et al., 2015; Lane et al., 2020).

Recently, wind energy projects are emerging in the Mediterranean Sea (International Energy Agency, 2019), an area within which information on potential wind farm impacts on seabird populations is virtually absent. This is concerning since the Mediterranean Sea is a biodiversity hotspot under immense pressure from cumulative threats (Micheli et al., 2013), especially for seabirds (Coll et al., 2012; Genovart et al., 2016; Genovart et al., 2018; Grémillet et al., 2018). We focused on the Scopoli’s shearwater, one of the three endemic shearwater species in the Mediterranean. The dark fate of French Scopoli’s shearwater populations may be accelerated by the race for decarbonated energy and planned offshore wind farm deployment along the French Mediterranean coast, notably with the deployment of an offshore wind farm within 50 km from the largest French shearwater colony (Ministère de la Transition Ecologique, 2022). We measured flight heights using highly precise barometric data loggers to anticipate the potential impacts of forthcoming offshore wind farms in the Gulf of Lion, on the population dynamic of Scopoli’s shearwater breeding in the Calanques National Park off Marseille, France.

## MATERIAL AND METHODS

### Barometric measurements

To estimate the flight altitude of Scopoli’s shearwaters, we equipped 13 breeding adults during the chick-rearing period (30^th^ July to 16^th^ August) with a barometer (MSR-145, MSR Electronics, Switzerland, 18×14×6 mm, 18 g) combined with a temperature-depth recorder with a wet/dry sensor (TDR, G5 CEFAS Technologies, UK, 7 mm diameter, 30 mm length, 2.7 g) in 2019 and 2020. We caught breeding adults by hand at the nest during nighttime. We attached barometers to back feathers with black Tesa® tape, and TDRs to plastic leg rings with black Tesa® tape. We recovered them at the nest after one to several foraging trips. Barometers recorded atmospheric pressure and temperature every second, whereas TDRs distinguished the periods when the bird dived or stayed at the sea surface (sensor was wet) from flight periods (sensor was dry). Tagging load represented ∼3.4% of average bird body mass (mean body mass in 2019-2020 = 640 g, range: 540 –765 g). Handling was generally performed in <15 minutes.

### Estimation of flight altitude

We adapted the framework of Cleasby et al. (2015) to estimate flight altitude of birds from pressure measurements for each flight bout. A flight bout was preceded and followed by a water landing phase, distinguished using the wet/dry state of the TDR loggers. We first decreased noise in the raw time-series of pressure data and smoothed pressure data with a running median calculated through a moving window with a range of seven seconds. We then converted pressure measurements into an estimation of flight altitude using the barometric formula (Berberan-Santos et al., 1997):

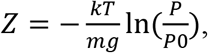

where Z is flight altitude in meters, P and P0 are atmospheric pressures in Pascal at altitude Z and at sea level, respectively, k is the universal gas constant for air (8.31432 N.m.mol^-1^.K^-1^), T is the temperature in degree Celsius of the atmospheric layer between sea level and Z, m is the molar mass of air (0.0289644 kg.mol^-1^), and g is the gravitational acceleration (9.80665 m.s^-2^). P and T were directly measured every second by the barometer. For each flight bout, we estimated P0 at each second as the mean between P0start and P0end (the mean of pressure measurements recorded in the final/first 10 seconds of the landing period preceding/following the current flight bout). When the landing period was < 10 seconds, we used its total duration. To consider temporal changes in the relative importance of P0start and P0final measures, we weighted P0start by the inverse of the square of the time elapsed since the start of the flight bout, and P0final by the inverse of the square of the time remaining until the end of the flight bout. Potential local changes in atmospheric pressure at the same location or displacements of birds towards areas with different atmospheric pressure at sea level bias P0 estimates.

Therefore, we only retained flight height estimates for which we had P0 estimates based on a measure of P0start and/or P0final less than 10 min old, the validity duration for P0 estimates found in Cleasby et al. (2015). We also retained only flight bout with a duration > 5 sec to avoid bias in pressure measures because of the strong turbulence during the take-off and landing. Overall, barometric flight height was estimated from 1934 flight bouts. Using the same pressure logger, Cleasby et al. (2015) showed that the mean error in flight height estimates of a seabird was 0.88 m (range 0.32-1.92 m).

## RESULTS

Using information from temperature-depth recorders (TDR, N = 13 birds), we found that birds at sea spent 71% of their time in contact with water and 29% in the air. Individual Scopoli’s shearwaters flew at very low altitudes with a mean height of 1.8 ± 2.7 m (± SD) above sea level (Fig. 1). Accordingly, shearwaters flew very rarely (<0.02%) within the vertical envelop of collision risk with wind turbines, i.e. above 20 m.

**Figure 1.**
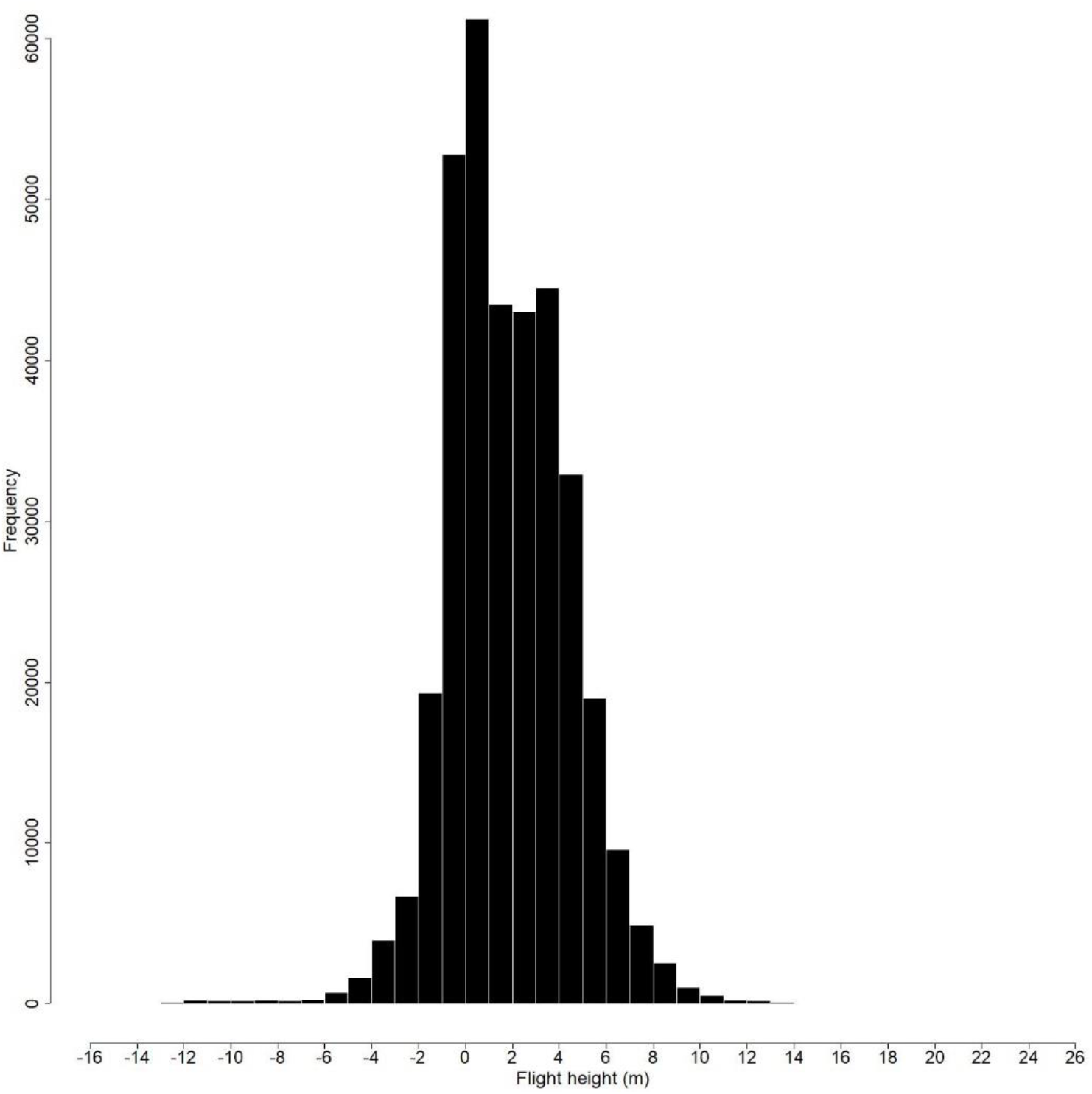
Density of Scopoli’s shearwater flight heights estimated from pressure measures.

## DISCUSSION

Our study provides the first flight height estimation for Scopoli’s shearwaters based on electronic recordings (independent of time of the day and weather conditions). It is of primary importance in the context of planned offshore wind farm deployment along the French Mediterranean coast close to the largest French shearwater colony (Ministère de la Transition Ecologique, 2022). Shearwaters usually fly ca. 1.8 m above the ocean surface. Almost half of the distribution had negative values, probably due to measurement errors and overpressure on the logger in certain flights circumstances near the sea surface. Overall, our flight height estimation is coherent with field observations and the fact that this species, as all petrels, uses dynamic soaring, a flight mode requiring proximity to the ocean’s surface (Cook et al., 2012; Deakin et al., 2022). We found that adult shearwaters rarely flew within the rotor-swept envelop (<0.02%), as reported for a closely related species, the Manx shearwater (*Puffinus puffinus)* in the Atlantic (0.04%, Cook et al., 2012). Therefore, we found that Scopoli’s shearwaters did not fly at collision-risk altitude and considered that collisions were probably negligible for population dynamic of Scopoli’s shearwater population from Marseille.

Yet, collision risk may increase in the future following behavioural modifications. Indeed, some shearwaters can be attracted to offshore structures due to local prey enhancement and/or light attraction, and are disoriented once attracted in the vicinity of a light source (review in Deakin et al., 2022). In addition to this potential but unknown collision risk, foraging habitat loss induced by planned wind farms (Peschko et al., 2021), a primary concern for seabird conservation (Dias et al., 2019), may also impact population dynamics.

Indeed, we found that birds routinely used the future area dedicated to wind farms in the eastern part of the Gulf of Lion to forage and rest (Courbin et al., 2018a). Loss of such functional habitat may compromise foraging strategy of individuals, with cascading negative effects on body condition, reproductive success and adult survival, as observed for Cape gannets *Morus capensis* in an overfishing context (Cohen et al., 2014).

## ACKNOWLEDGMENTS

This study was funded by the Ademe and energy firms (EDF Renouvelables, Engie, Eolfi) within the program ORNIT-EOF, the French Office for Biodiversity within the program INDEXPUF, and by the OSU OREME Montpellier. Thanks to Alain Mante and the Parc National des Calanques. Thanks to G. Herrouin from Pôle Mer Méditerranée for coordinating ORNIT-EOF program, and all fieldworkers involved in this study. Handling protocols for shearwaters were approved by the boards of the Calanques National Park (permit number: 2016-111), the French ‘Direction Départementale de la Protection des Populations’ (permit number: 34-369, #A34-505), the French ‘Direction Départementale des Territoires et de la Mer des Bouches-du-Rhône’ (permit number: 13-2019-07-25-011) and the ‘Comité d’Ethique pour l’Expérimentation Animale Languedoc-Roussillon’ (permit number: 1170, APAFIS#20842-2019052710179096).

## Notes

### Competing Interest Statement

The authors have declared no competing interest.

